# Progressive Backmapping of Highly Coarse-Grained Protein Models

**DOI:** 10.64898/2026.03.02.709104

**Authors:** Yu Zhu, Jacob M. Remington, Shenghan Song, Bo Yang, Brody P. Magee, Severin T. Schneebeli, Jianing Li

## Abstract

Reconstructing all-atom (AA) structures from highly coarse-grained (HCG) models remains a significant challenge in multiscale molecular dynamics (MD) simulations, particularly for mesoscale biomolecular assemblies that are beyond the reach of conventional MD methods. Building upon ProNet Backmapping, a neural-network–based thermodynamically consistent approach, we introduce a progressive backmapping framework that reconstructs AA models in a stepwise manner across neighboring resolutions, for example, from a 3-residue-per-site HCG model to a 1-residue-per-site model, then to an AA model. This progressive backmapping method achieves high accuracy across a wide range of proteins and effectively reconstructs flexible linkers in multidomain architectures. Moreover, it supports hierarchical reconstruction of complex protein assemblies, including multiple virus-like particles spanning tens of nanometers and containing hundreds of subunits. Using this framework, we demonstrate—for the first time—the ability to hierarchically backmap entire viral assemblies from HCG to full AA resolution, covering at least three different resolutions. Overall, our method provides a scalable framework for incorporating atomistic detail into mesoscale simulations of complex systems across many applications in chemistry and biology.

**Table of contents figure:** 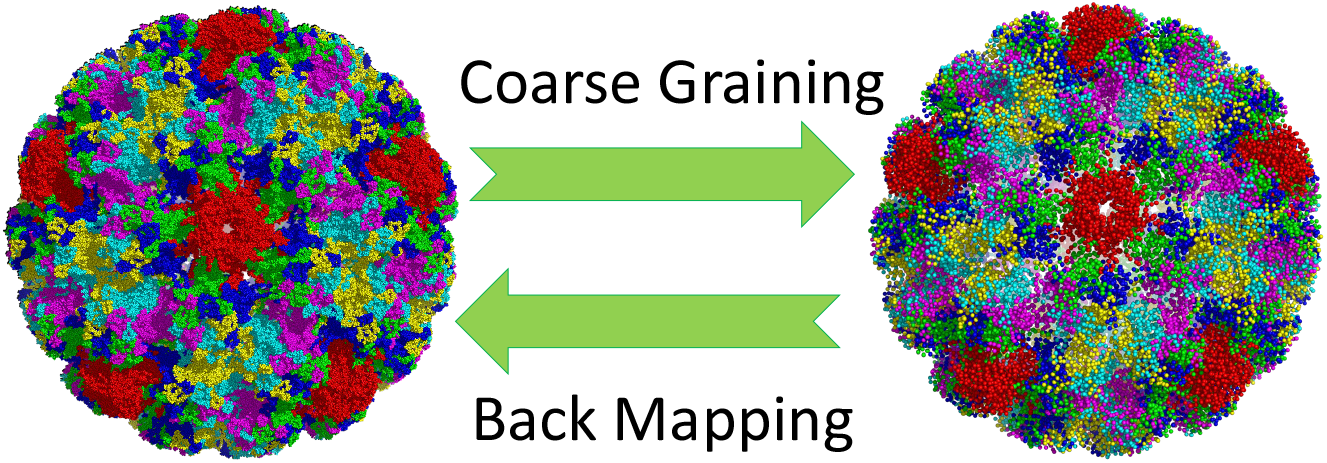

## Introduction

The ability to computationally examine the structures and dynamics of biological macromolecules and their complexes across various time and length scales is essential for many applications in chemistry and biology. These applications include understanding and predicting self-assembly, aggregation, and host-guest binding, which are vital to design new materials and medicines. Despite considerable progress in multiscale modeling,^1–6^ current structure-based modeling techniques still face significant limitations in obtaining essential atomic details on mesoscopic scales. Although coarse-grained (CG) models are commonly used to access mesoscopic time or length scales by representing macromolecules as linear combinations of atomic positions that preserve essential shape and features,^7–10^ atomistic insight remains essential for encoding biophysical properties and enabling biochemical functions, particularly for rational design purposes, like enzyme engineering and nanomedicine design.^11–13^ Prior studies by us and others suggest that combining CG and all-atom (AA) models can be useful for studying a wide range of proteins and protein assemblies in different environments, which cannot be achieved with a single model resolution.^14–19^ To overcome this fundamental challenge, an AA model can be transformed into multiple CG resolutions, with some closer to the AA resolution (such as the UNIRES, MARTINI-CG, and SIRAH models)^4,7,20–24^ and others much coarser (such as ultra or highly coarse-grained models) within hierarchical models.^10,25–27^ Nevertheless, how to systematically integrate protein models across varying resolutions and obtain essential atomistic insights at mesoscopic scales still remains unresolved, especially as it relates to accurately recovering the AA details from a highly coarse-grained (HCG) molecular dynamics (MD) simulation.^1^

Herein, we present a general, progressive backmapping framework that uses an AI-/ML-guided method to effectively recover the coordinates of the underlying, more detailed representations within hierarchical models. Using three model resolutions (MARTINI-CG, AACG, and AA), we tested both a bottom-up approach for membrane protein dimerization^28^ and a top-down approach for peptide self-assembly,^14^ demonstrating proof of concept for hierarchical modeling, including successful hierarchical backmapping from relatively finer-grained models to AA models. Coarser models like HCG models are generally required to significantly accelerate sampling, with notable examples including the resolution replica exchange method developed for exploring complex energy landscapes,^29^ backmapping biased sampling (BMBS) to enable sampling on chemically and biologically relevant time scales,^30^ and top-down multiscale approaches for studying the SARS-CoV-2 virion^31^. Yet, successful hierarchical modeling and especially atomistic detail recovery from HCG models have not been shown previously. In general, the core of hierarchical modeling and simulation is the accurate interconversion, where transformations from higher to lower-resolution models (known as mapping) and from lower to higher-resolution models (known as backmapping) are both essential. Notably, there is a particular need for more effective hierarchical mapping and backmapping methods in nanomedicine. With the rapid advancement of nanomedicine in recent years, MD simulations of large, complex protein assemblies have become useful for modeling their mesoscopic properties. These assemblies consist of hundreds of protein subunits arranged in highly symmetric architectures, such as adeno-associated viruses (AAVs) for gene therapy delivery^32^ and virus-like particles (VLPs) for vaccine design.^33^ In such applications, AA resolution is indispensable for probing stability, toxicity, and immunogenicity, yet only HCG models—coarser than one residue per site—can feasibly access mesoscopic length and time scales. Bridging this resolution gap demands accurate mapping and backmapping strategies that integrate protein models hierarchically to enable realistic AAV and VLP design.

Currently, mapping from AA to CG can be achieved via several approaches, such as thermodynamic fluctuations,^34,35^ topology and graph theory,^7,8^ or even intuition.^36–39^ However, backmapping from CG to AA protein models, particularly from HCG to AA, is considerably more challenging for several physical reasons. First, while specialized backmapping methods have been developed for small molecules,^40–42^ polymers,^43,44^ and lipids,^45–47^ they have limited transferability and cannot be readily generalized to biomolecules such as proteins. For example, a recent work from Pezeshkian *et al*. introduced a multiscale algorithm that can backmapping triangulated lipid membrane surface into their corresponding MARTINI-CG representations.^45^ While this approach performs exceptionally well for lipid membranes, it cannot be directly applied to proteins because of their unique structural features and the finer resolution typically used in protein models. Second, as the CG models can span a broad range of resolutions, the difficulty increases with the degree of coarse graining because less information is retained for backmapping, especially in HCG models. Most existing protein backmapping approaches are designed to reconstruct AA models from CG representations with resolution finer than one site per residue. For example, Bennett *et al*. showed that their ezAlgin can backmapping various MARTINI-CG models, including lipids and transmembrane proteins, to AA structures^48^. Some methods support one site per residue. For instance, Lombardi et al. presented a CG2AA scheme capable of reconstructing AA structures from any model containing beads at the alpha-carbon (C_α_) atom. However, few methods are suitable for backmapping from HCG to AA models, because achieving high accuracy is more difficult due to the limited information retained in HCG models, unlike CG models such as MARTINI that operate at finer resolutions than one site per residue. Therefore, developing a general backmapping method that accurately converts HCG models into AA structures for general protein systems is urgently needed.

The methods of backmapping can be categorized into two directions: naive rule-based approaches and data-driven approaches. Early attempts predominantly relied on rule-based methods, including backmapping multi-resolution CG Models to the AA model using Bayesian inference and constrained MD simulations,^49^ reconstructing protein backbone and sidechain separately by combining both geometrical information with the MARTINI mapping scheme,^50^ and replacing CG sites with predefined AA fragments by inquiring a database of fragment files. These rule-based approaches typically depend on predefined CG mapping rules, geometric relationships, a specific dataset, or energy minimization procedures. However, due to the inherent degeneracy of CG mapping, such approaches often struggle when explicit rules are unavailable, tend to be overly deterministic, fail to capture thermodynamic information across diverse CG conformations, and are typically limited to resolutions finer than one site per residue.^24,49,50^ As CG resolution becomes coarser and information content decreases, recovering the high-dimensional atomic configuration of proteins becomes extremely challenging. Consequently, leveraging global variables, such as radius of gyration^51^ and interdomain center of mass (COM) distances^52^ to local variables like secondary structure^53^, sidechain torsions^54^, or contacts^55^ becomes essential, as these key features can be preserved in a surprisingly small number of degrees of freedom, such as HCG models.

To address the limitations of rule-based methods and better incorporate these global variables, several data-driven techniques have been developed in recent years by utilizing machine learning technology, including Variational AutoEncoders (VAEs),^41,56^ and Generative Adversarial Networks (GANs).^42,43^ While these methods demonstrate strong performance, many have been applied to polymers^43^ and small molecules.^40^ For example, Li *et al*. employed a conditional GAN to backmapping polymer CG models,^43^ while An and Deshmukh used four regression neural network models to backmapping CG carbon chains.^40^ However, due to the substantial structural differences between these systems and proteins, dataset selection and training costs make it difficult to directly extend such approaches to protein backmapping. Using E(t)-transformer model, Mahmoud *et al*. have shown that it can backmapping high-resolution SIRAH CG (finer than 1 residue per site) representation of proteins to AA structures.^57^ More recently, Jones *et al*. introduced a diffusion-denoising autoregressive model for the non-deterministic backmapping method to restore AA details from C_α_ coordinates.^58^ Their method reconstructs proteins sequentially from the N-terminus to the C-terminus, residue by residue, conditioned on the C_α_ trace and local environment. The model takes a Cartesian coordinate representation of the target residue, the local environment, and one-hot encoding of the residue identity as the input. Because this non-deterministic method is designed to generate a diverse ensemble of AA structures, the predicted structures are not optimized to minimize deviation from the reference. Consequently, its applicability is limited to tracking continuous trajectories and extending to HCG models, which require high alignment accuracy.

In contrast, this work now establishes a progressive backmapping approach that performs the conversion stepwise across neighboring resolutions. Our previous work^8^ on the Fixed Length Coarse Graining (FLCG) approach has demonstrated that HCG models with resolutions as coarse as 6 residues per site can still capture protein thermodynamic properties. FLCG defines the coarsest HCG resolution and develops the geometry-based rules for backmapping from HCG to the 1-residue-per-site CG resolution.^8^ Rather than backmapping directly from HCG models to AA models, a given protein system can be first built as a 6-residue-per-site HCG model, then as a 3-residue-per-site HCG model, and finally as a 1-residue-per-site CG model, based on these simple geometry-based rules.^8^ From the 1-residue-per-site CG model, in this work we have developed ProNet Backmapping, a neural network-based position-matching method to construct the thermodynamically consistent AA model. Notably, progressive backmapping of large-scale protein complexes is particularly challenging because errors accumulate from coarser to finer-grained models. Most existing approaches begin from C_α_-only representations when reconstructing AA structures,^50,58^ which often limits the achievable agreement between predicted and reference structures. To our knowledge, this is the first demonstration of accurate HCG backmapping capabilities for general protein systems and biomedical applications.

In the following sections, we first present the theoretical foundation and implementation details of the ProNet Backmapping approach and then report prediction results across a wide range of protein lengths and structures for both the training and validation datasets. We also demonstrate the applications of ProNet Backmapping to flexible proteins, such as multidomain proteins (MDPs), and then show progressive backmapping on large protein assemblies, including adeno-associated virus 2 (AAV2) and human papillomavirus (HPV). Compared with existing methods, our backmapping approach achieves better agreements between reconstructed and reference structures, delivering state-of-the-art performance in several key aspects: (1) accurate reproduction of the thermodynamic properties; (2) high-fidelity reconstruction of flexible complex proteins; (3) general applications spanning from simple proteins to large assemblies; (4) novel applications to study protein mutants and variance. A summary and conclusion are provided at the end to highlight the future applications.

### Theory & Methods

#### Position Matching and Thermodynamically Consistent Backmapping

The fundamental assumption of hierarchical modeling is that simulations performed with models at different resolutions, such as HCG, CG, or AA models, should yield consistent results. This assumption implies that a backmapped AA model should also be consistent with the original CG representations. Consider a molecular system described at two different resolutions: an AA atom model and a CG model. The AA model has higher resolution and contains more particles (*n*) than the number of sites (*N*) in the CG system. To ensure consistency between two representations, previous work by Noid *et al*. have shown that two key constraints must be satisfied^9^: (1) The coordinates and momenta of each CG site must be assigned by a well-defined linear combination of coordinates and momenta of a subset of the atoms from the AA systems (2) The equilibrium distribution of coordinates and momenta of CG sites must match between both AA and CG models. In practice, however, many CG models treat momenta implicitly^59^ or apply geometry-based rules^36^ to CG models, which only ensure that the equilibrium distribution of CG coordinates aligns with that in AA models. As a result, maintaining consistency in the coordinate representations between the CG and AA systems becomes more critical than matching momenta.

For an AA system consisting of *n* particles with position denoted as **r**^*n*^ = {**r**_1_ … **r**_*n*_}, and a CG system with *N* sites and their position given by **R**^*N*^ = {**R**_1_ … **R**_*N*_}, the equilibrium distribution of coordinates of both systems are described by the following two equations, where *u*(**r**^*n*^) represents the potential energy of the AA system, and *U*(**R**^*N*^) is the potential energy of the CG system:

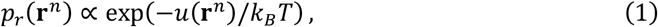

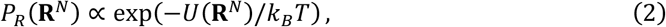

To ensure consistency between the CG model and the AA model in configuration space, the following relationship between their distributions must be satisfied,

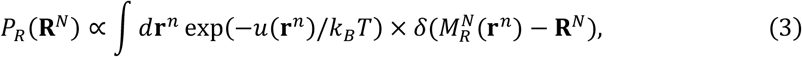

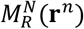 is a specific linear mapping operator to covert the coordinates in atomistic system into coordinates of CG sites in CG system, In the following context, 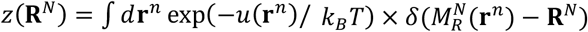, for simplify.

In backmapping, the conditional probability distribution, *P*(**r**^*n*^|**R**^*N*^), which is used to reconstruct the AA structure from a given a certain CG configuration, can be derived by applying Bayes’ Theorem^60^ to Eq.3.

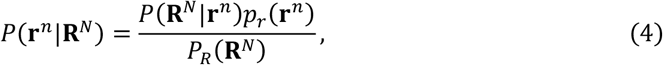

Such that,

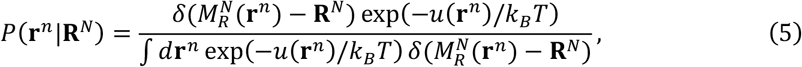

This probability function applies to any continuous quantities that can be derived from AA coordinates.^9^ The delta function term, 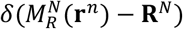, ensures that the probability density is non-zero density only when 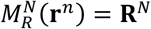, which enforces that the backmapped AA coordinates match the given CG configuration, and lie in the 3*n* × 3*N* dimensional subspace, denoted as 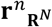, which contains AA configurations consistent with the same CG mapping. The Boltzmann term, exp(−*u*(**r**^*n*^)/*kBT*) weights atomistic structures by energy. The normalization factor, (1/*z*(**R**^*N*^)), acts as a partition function, providing a proper measure over the probability space. As a result, the AA structure with relatively lower energy within the subspace 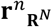 are more likely to be generated.

Since Eq.5 defines the probability of an AA configuration given a CG model, it effectively quantifies the degree of consistency between the two resolutions. This formulation provides the theoretical foundation for constructing a thermodynamically consistent^9,10^ backmapping model. In contrast to other deterministic models or predefined mapping approaches that rely on a direct mapping function to produce a single AA structure from a CG input, this approach can maintain thermodynamic consistency by allowing the mapped subspace to contain an ensemble of AA configurations rather than a single output defined by the function.

While a deterministic model may not generate diverse samples from *P*(**r**^*n*^|**R**^*N*^), an appealing approximation is to select structure where *P*(**r**^*n*^|**R**^*N*^) is maximized. This could yield better structural alignment between the predicted AA structures and the reference configurations compared to non-deterministic models that draw structures randomly from the probability distribution. However, the maximization process is numerically challenging for this high-dimensional problem. Alternatively, the ensemble-average structure, 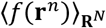, given by Eq. 6 is appealing because it is mathematically similar to the force-matching theory^61^ and the methods used to determine the CG potential energy function.^9,10^

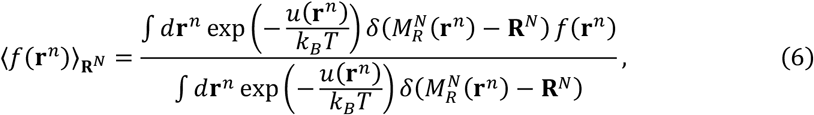

Namely, the variational principle for force matching also suggest that an optimal model for 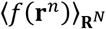 can be determined by position-matching atomistic structures sample from canonical ensemble. In the following paragraph, we parameterize such a model using an artificial neural network and demonstrate sampling of 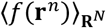 for any protein system containing the canonical amino acids.

Developing a deterministic backmapping approach for proteins using position-matching presents a multitude of practical challenges. Proteins are often structurally flexible, and both the combinatorics of protein sequence and the sampling problem can preclude complete convergence of 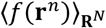 of any specific protein. For convenience, we decompose the problem into predicting each residue’s structure from a set of features instead of predicting the entire protein in one shot. The primary features are the one-hot encoding of the CG site amino acid, and a featurization of CG site’s environment as follows. Following the principle of locality, the position of a single bead of a structure, is influenced directly by its surroundings, we first include the *k*-nearest neighbors, R_*i,k*_ as the input to help in convergence, where *i* the index of the residue of the protein, and *k* is the number of the neighbor residues. Previous works shown that the neural network models to predict energy or forces from distances along facing several disadvantages^62,63^, due to that (1) the coordinates of configurations are fed into the NN sequentially; (2) the interchange of two atom’s coordinates will change the total energy in the neural network model. Thus, the three-dimensional structure is encoded in the form of special atom-center descriptors, and the distance between our CG sites are transformed using weighted radial symmetry functions^62^, rather than directly using cartesian coordinates, which is constructed as a sum of Gaussian with the scale and center parameters *η* and *r*_*s*_.

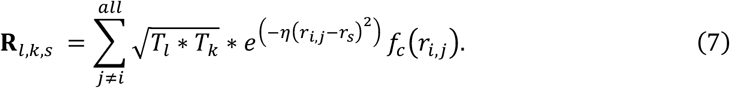

A key hypothesis in Eq. 7 is that the 20 amino acids can be further generalized into a smaller number of bead types, *L*, (*L*=4 used in this work) and the relative locations of beads and their types is the main driver for intra-bead atomic positions. For example, when backmapping a positively charged ARG, it’s side chain conformation will be heavily influenced by the location of neighboring negatively charged residues, be it ASP or GLU. Since the number of interactions pairs is *L*(*L* + 1)/2, computational expense can be saved by using a small L. The subscript *l* and *k* indicate the type any two different CG sites and the rescaling term given by *T*_*l*_ and *T*_*k*_ are weights^64^ derived during training by a CG type neural network mapping one-hot encoded amino acids to vectors of length *L*. Finally, a cutoff function 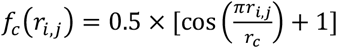 is used to define the energetically relevant local environment. This function only works when the distance *r*_*i,j*_ between two CG site smaller than cutoff *r*_*c*_, and yield zero when *r*_*i,j*_ > *r*_*c*_. The values of **R**_*l,k,s*_ are used as features that are fed into the neural network. To help in convergence, we also include local structure of the CG tripeptides from the neighboring two bonds and one angle of triplet into the features.

Another challenge arises from the fact that the rotational and translational invariance (RTI) of *u*(**r**^*n*^) implies that the ensemble average in 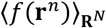 must be taken in an internal reference frame or with internal coordinates. To enforce RTI in our position matching procedure, we first generate 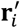 in the local frame of the protein chain defined by the CG positions of three sequential amino acids (R_*i*−1_, R_*i*_, R_*i*+1_ for interior residues). Predictions in the local frame are then aligned to the protein chain on-the-fly during training using root mean squared displacement (RMSD) alignment of the CG sites and Kabsch algorithms^65^ as described in Eq (8), where *T*_*i*_ is the transformation matrix.

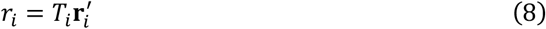

In addition, in Eq.5, the term 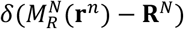 indicate that atomistic structure need to lie in the subspace 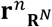, and only happened when 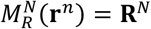. This is important when considering that successive backmapping and CG mapping should reconstruct the same CG structure. To apply this restriction, we use an orthogonal basis for subspace 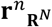, and using liner combinations of atomic positions, that are orthogonal to rows of 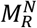, defined as 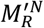, to ensure the predicted positions in line with subspace 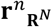. So, before the coordinates 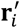 aligned using Eq (8), another transformation is required using Eq (9).

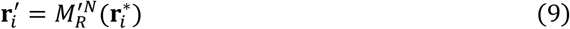

Finally, since the ensemble average structure may come with relatively higher energy, which is mainly due to torsional rotations, a quick energy minimization using *Amber*^66^ software is performed in the last step to fine tune the prediction.

#### Large Dataset for Training and Validation

The dataset is preprocessed (ESI Figure.S1) and constructed using AA molecular dynamics simulations of 320 proteins (ESI Table.S1) selected from the Protein Data Bank (PDB) and totaling 50 *ns* stored with 10,000 frames were used to approximate the CG environment in the canonical ensemble. 2.5 million frames from AA simulations of 250 proteins were randomly selected for model training, and 0.7 million frames from 70 structurally distinct proteins were used for validation to demonstrate broad generalizability. The *pdb4amber* command was used to remove hydrogen and water in.*pdb* files, and then systems were solvated with optimal point charge water^67^, neutralized with the requisite number of Na^+^ or Cl^-^. All proteins are simulated using force filed ff19SB^68^ with a Langevin thermostat^69^, and executed at the temperature 300K using *Amber*^66^ software. The length of proteins in our dataset across a wide range, varies between 200 and 2400 residues (see Figure.1(B)), and the selection of proteins is not limited by any specific conditions related to their functions, sizes, or other characteristics. The CG coordinates are obtained from the center of geometry of atoms within each residue, which is described by Eq (10), where *m* is number of atoms within *i*_*th*_ residue,

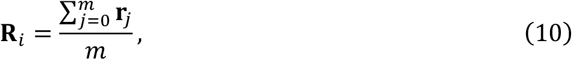

#### Implementation Details of ProNet Backmapping

To facilitate the training of the ProNet Backmapping approach, the workflow of data preparation is illustrated in Figure 1A. First, the PDB structures in our dataset were prepared and simulated using the AMBER software package. Second, the protein in each frame was coarse-grained into a 1-resisde per site CG representation. Third, structural features, including coordinates, bonds, angles, and neighboring environment, were extracted and vectorized as model input. The architecture of the ProNet Backmapping approach is shown in Figure 1D. In the top panel, the identities of CG sites of 3 site segments and their nearest neighbors are represented as a fixed-length fingerprint using a one-hot encoder. These fingerprints are then transformed into a dynamic fingerprint through three regular densely connected layers, as dynamic fingerprints have been shown to provide better predictive performance.^70^ The coordinates of the triplet segment and its nearest neighbors, together with the encoded identities, are transformed into **R**_*l,k,s*_ using the weighted radial symmetry function, Eq (7). Subsequently, the bond, angle, and identity features of tripeptides are processed through four fully connected neural networks. The decoded positional features from the fourth layer are transformed using Eqs (8) and (9) to predict the AA coordinates. The network includes three hidden layers with reduced dimensionality in the Identities encoder to encode residue identities, and four hidden layers in the positions decoder are implemented for parameter optimization. A leaky rectified linear unit (Leaky ReLU) activation function is used throughout, with a negative slope coefficient of 0.3. L2 regularization is applied with a coefficient of 0.01. The Adam optimizer is employed with a learning rate of 0.00025. The loss function is defined by Eq. 11 which measures the discrepancy between the reference AA coordinates and the aligned predicted coordinates obtained using Eq. (8),

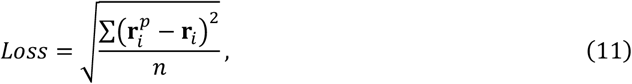

Where 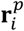 represented the coordinates of predicted atomistic structure.

**Figure 1.**
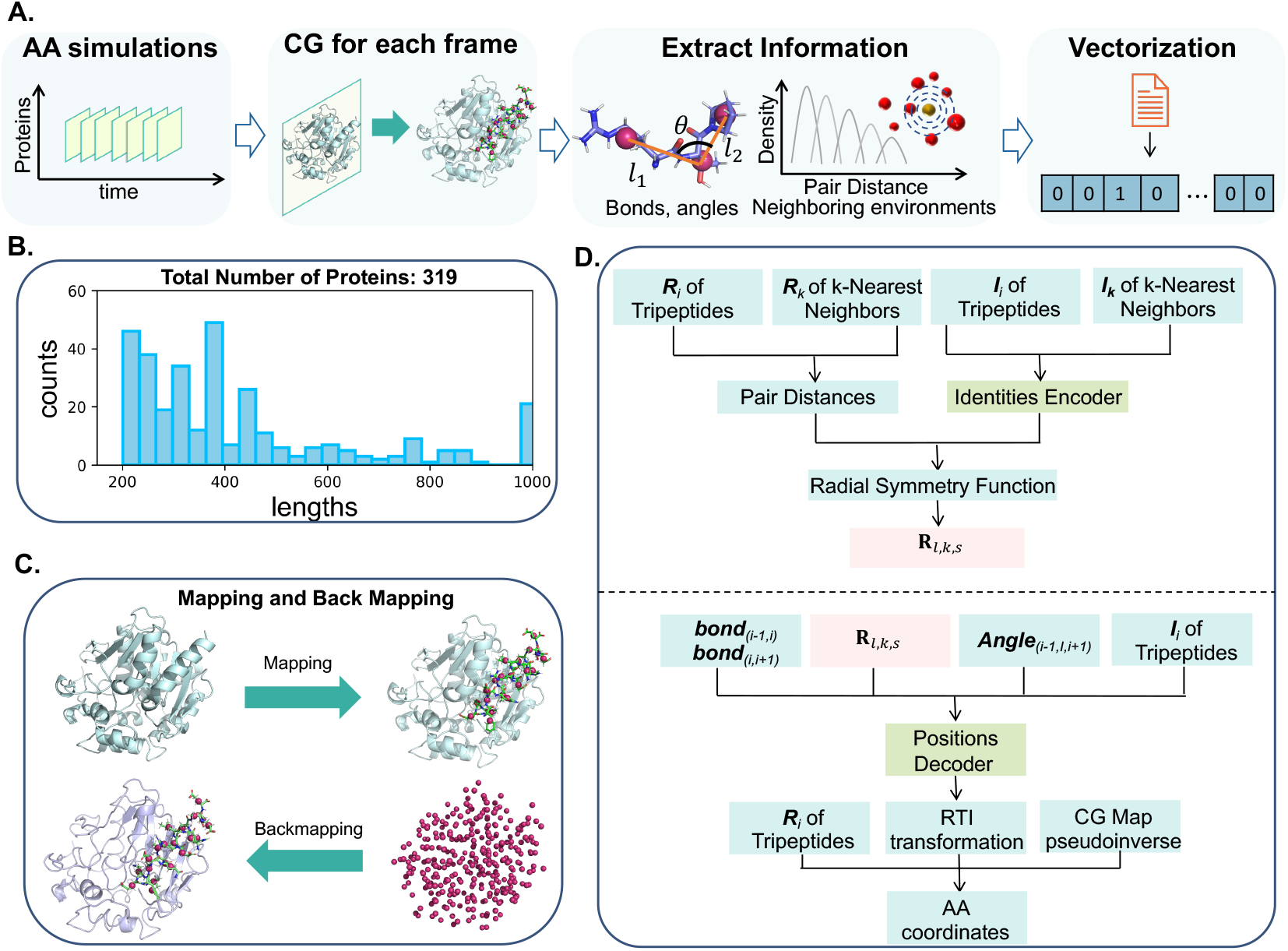
(A) Workflow of data processing and feature extraction. (B) Distribution of protein length (in units of residue) in our dataset. (C) The figure depicts the coarse-graining process that converts the residues(green) into CG sites (red), where 1 bead represents a residue; and a list of CG sites can be backmapped into AA structure residue by residue. A portion of the protein is represented as sticks together with CG sites to illustrate their alignment. (D) The architecture of ProNet Backmapping model for atomistic model prediction. The top panel presents the encoder process of **R**_*l,k,s*_ using Eq. 7 while the bottom panel presents the decoder process for coordinates aligned with rotational and translational invariance transformation. The identity encoder and position decoder units are constructed using neural network models.

Figure. 1C illustrates the coarse-graining process, in which a detailed AA structure is simplified into a set of representative sites by computing their COMs, as well as the backmapping process, where these sites are reconstructed into the original AA protein structure using our well-developed backmapping algorithm.

#### Characterization of Backmapping Agreement

The agreement between the backmapped and reference structures are characterized using root mean square fluctuation (RMSF), calculated according to Eq. (11), where *x*_*i*_ is the coordinates of residue *i* or CG site *i*, ⟨*x*_*i*_⟩ is the ensemble average position of *i*.

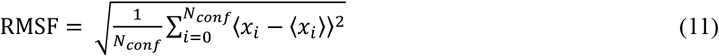

A small value indicates that each residue in the backmapped structures is well aligned with the reference structures.

## Results and discussion

### Evaluation of Predictive Performance

The backmapped or reconstructed AA models generated by ProNet Backmapping maintain consistently high structural fidelity across a wide range of proteins in both training and validation datasets. They display a narrow RMSD distribution centered between 1.6 and 1.7 Å against reference X-ray or cryo-EM structures (Figure 2A). The similar distributions seen in both the training and validation sets indicate high predictive accuracy and good generalization. Importantly, reconstruction quality remains stable across proteins of varying sizes and structural motifs. The RMSD metric shows minimal dependence on protein length (Figure 2B), indicating that ProNet Backmapping scales robustly regardless of protein sizes. The backbone RMSD values are low, averaging at 1.2 Å, while the sidechain RMSD values remain below 2 Å, which suggests our approach well capture features in the local sidechain and global folding of proteins. To further illustrate these trends, we selected 20 proteins from training set and 20 proteins from validation datasets at uniform intervals after sorting by RMSD values in ascending order (Figure 2C and 2D).

**Figure 2.**
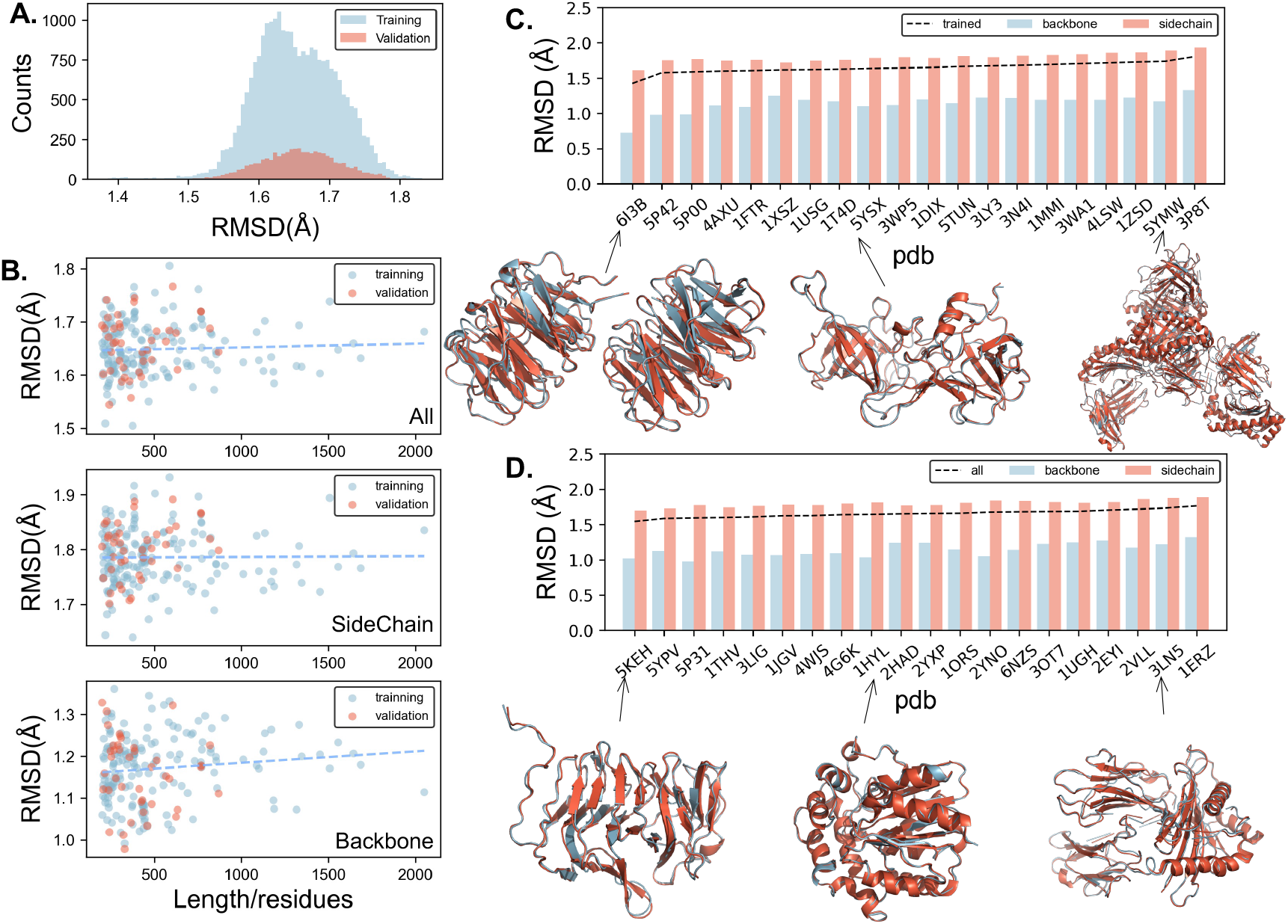
(A) Distribution of overall RMSD across the training and validation datasets. (B) Overall, backbone, and sidechain RMSD as functions of protein length. RMSD for 40 representative proteins from (C) the training dataset and (D) the validation dataset. The snapshots show proteins (cyan) backmapped from their 1-residue/site CG models in both training and validation datasets. The orange represents in snapshots are obtained experimentally.

Across all 40 representative proteins, the backbone RMSD is consistently lower than that of the side chains. Specifically, backbone RMSD values are approximately 1 Å, whereas sidechain RMSD values are close to 2 Å.

Representative cases highlight the high fidelity of the reconstructions. For the case of the truncated hemolysin A protein (PDB: 5KEH)^71^ ProNet Backmapping achieves an RMSD of 1.55 Å, with backmapped AA model in good agreement with the experimental structure, as shown in Figure 2. Notably, this RMSD is comparable to the 1.55 Å resolution of the reference PDB structure from X-ray diffraction,^71^ indicating that ProNet Backmapping can potentially achieve near-experimental accuracy. Although a few backmapped AA models show slightly higher RMSD values, these deviations still remain below 2 Å across both the training and validation datasets. Examples include the PDB structures 5YMW and 3LN5 shown in Figure 2, whose somewhat larger deviations can be attributed to their increased sequence lengths. Notably, the corresponding experimental reference structures—determined by X‐ray crystallography with reported resolutions of 2.0 Å and 1.9 Å, respectively—exhibit similar levels of inherent structural uncertainty. Thus, the magnitudes of the RMSD values for our backmapped models are comparable to the resolutions of the experimental structures. Overall, the AA models reconstructed by ProNet Backmapping accurately preserve global topology and maintain strong structural fidelity across proteins of varying sizes. Most of the discrepancies arise from sidechain atoms, whereas backbone RMSD values remain substantially lower. Together, these results demonstrate that ProNet Backmapping provides reliable, size‐independent reconstruction performance with near‐experimental accuracy.

Most protein structure backampping models are designed to predict a single rigid, static conformation corresponding to the most stable structural state. In contrast, our backmapping framework reconstructs AA structure from consecutive simulation frames, therefore directly bridging the MD simulations across different resolutions while substantially reducing computational cost. Crucially, this design enables the model to capture protein dynamics rather than only static structures. To evaluate this capability, we examined three representative proteins (PDB structures: 5KEH, 2HAD, and 3LN5) and assessed whether the reconstructed structures remain within the thermodynamics space. Across full AA MD trajectories, the RMSD of the backmapped models (orange lines) closely tracks that of the corresponding AA simulations (blue lines) over time (Figure 3A1-C1). As thermal fluctuations drive continuous conformational changes, the backmapped AA models faithfully track these transitions, indicating that the method accurately captures evolving structural states rather than collapsing into a single conformation.

**Figure 3.**
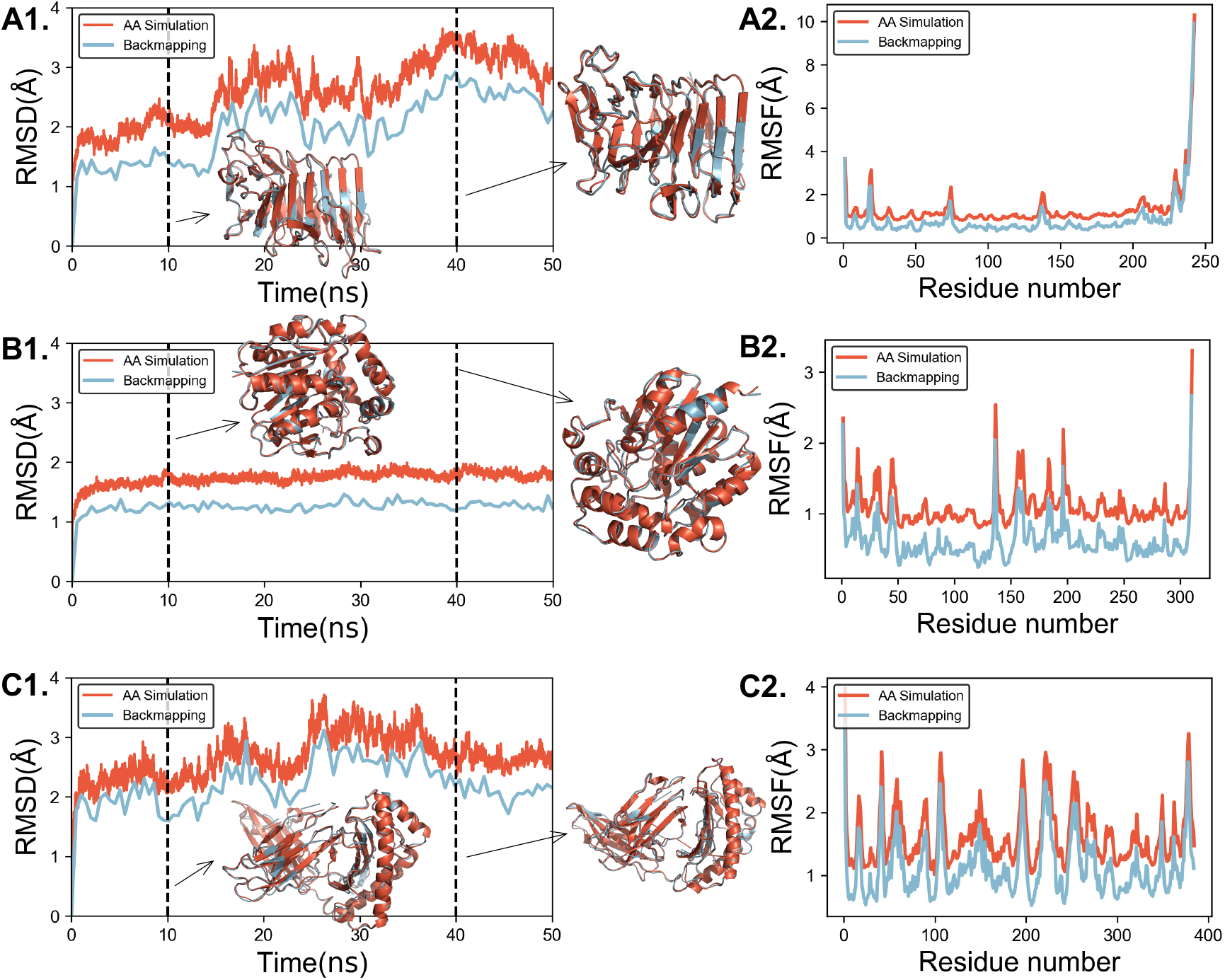
RMSD(A1-C1) and RMSF (A2-C2) of the AA simulations and backmapped AA structures within 50 ns for PDB structures 5KEH, 2HAD and 3LN5. RMSF values of the MD simulated structures were shifted upward by 0.5 Å for clarify. Snapshots of the backmapped (blue) and MD simulated (orange) structures at 10ns and 40 ns are shown for PDB structures 5KEH, 2HAD, and 3LN5.

Snapshot comparisons at 10 and 40 ns in Figure 3 further illustrate that the backmapped conformations remain well aligned with their AA counterparts, indicating that the model can reproduce multiple conformations consistent with the thermal ensembles. Moreover, RMSF analysis relative to trajectory-averaged structures shows strong agreement between AA simulations and backmapped models across all three proteins (Figure 3A2–C2). The consistent overlap of RMSF profiles confirms that residue-level flexibility and dynamic patterns are faithfully preserved. Unlike other predictive model, which typically output a single most probable structure, our approach generates ensembles of conformations that reflect the intrinsic thermal dynamics of proteins. These results demonstrate that ProtNet Backmapping not only reconstructs accurate static geometries but also maintains dynamic consistency across multiscale simulations.

### High-fidelity recovery of linker regions in multidomain (MDP) proteins

MDPs are ubiquitous, accounting for more than 80% eukaryotic proteins and approximately 67% of prokaryotic proteins. In these proteins, individual domain within an MDP typically evolves to fulfill a distinct function-such as catalysis, environmental sensing, host-guest recognition, or signal transductions. These domains are connected by short or long linkers and work together in a coordinated manner to enable complex physiological functions. Although predictive models can predict individual domains very accurately, especially if each domain is well represented in the training data, they often struggle with flexible linkers or dynamic inter-domain interactions. This limitation arises from uncertainty in relative domain-domain orientation and intrinsic flexibility between domains, making intrinsically disordered linkers especially challenging. In contrast, the backmapping method can yield improved performance on MDPs, as it inherently reconstructs atomic-level structures based on CG template. This temporal continuity allows for smoother transitions between conformations, preserving the dynamics interplay between domains, highly reduce the uncertainty in the disordered regions.

As shown in Figure 4, RMSF profiles of 6 representative MDPs exhibit strong agreement between AA simulations and backmapped models, and corresponding snapshots further illustrate the fidelity of the reconstructions. For each protein, four predicted AA models are presented, demonstrating that ProNet Backmapping approach accurately captures both the relative orientation between domains and their flexible linkers. This capability highlights the robustness of our method in reconstructing domain arrangements with high fidelity, even in the presence of conformational variability. Moreover, these examples along with additional MDPs (ESI Figure S2), as well as other small proteins (ESI Table S2) were not explicitly included in the training set. The consistent performance across these systems once again suggests that ProNet Backmapping approach generalizes well and can be reliably extended to more complex multi-domain systems where linker flexibility plays a crucial role in functions.

**Figure 4.**
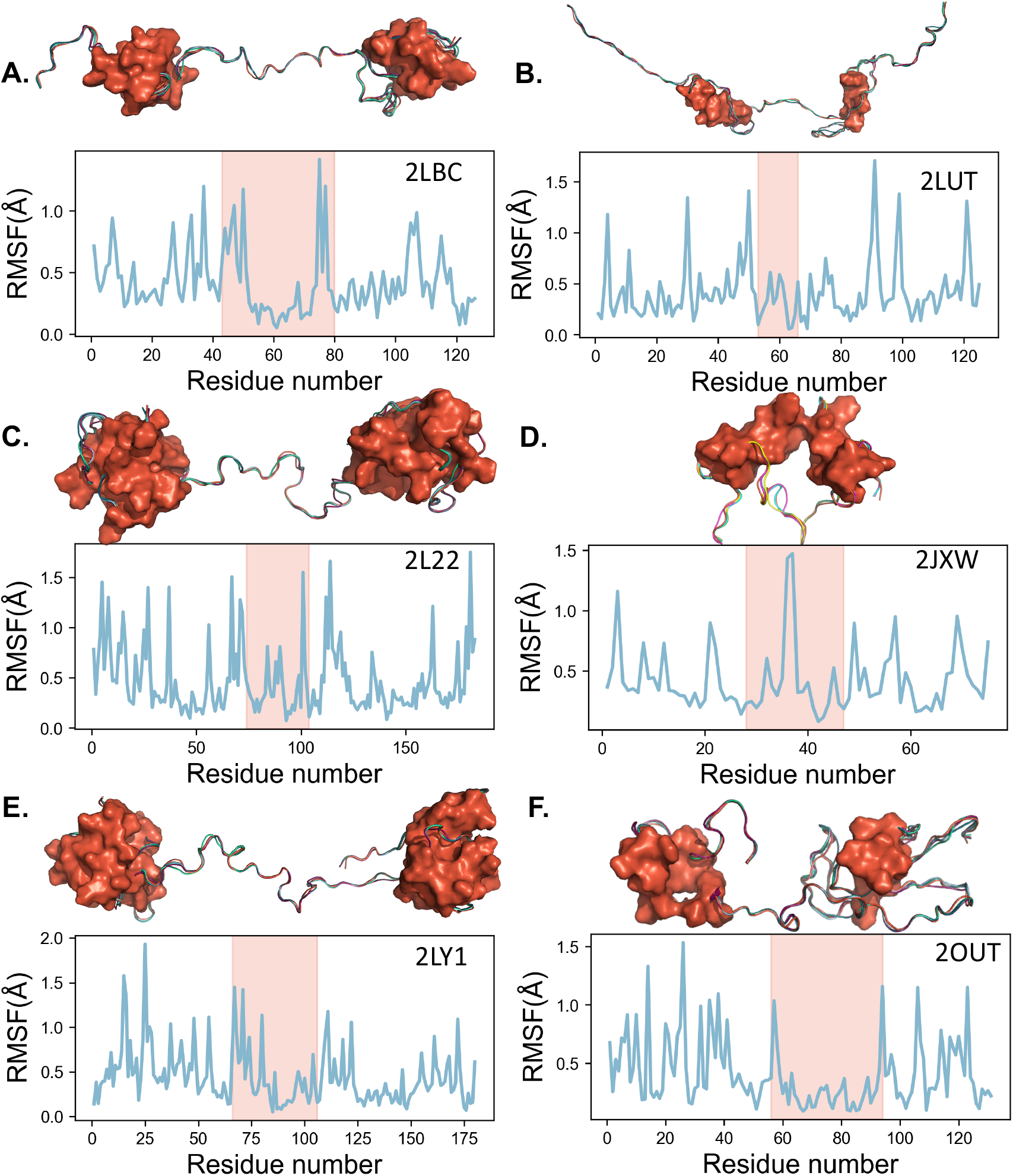
RMSF of the two-domain proteins, including PDB structures 2LBC (A), 2LUT (B), 2L22 (C), 2JXW(D), 2LY1 (E), and 2OUT (F), along with their corresponding snapshoots. The linker regions are highlighted in pink. In the snapshots, domain regions are shown as an orange surface, with each snapshot containing one experimentally obtained structure and four backmapped models.

### Progressive backmapping of protein assemblies from HCG to AA models

Backmapping is particularly valuable for large protein assemblies because AA simulations of these systems are computationally prohibitive. Viral capsids such as AAV and HPV contain tens to hundreds of protein subunits and span tens of nanometers in diameter. AAV2 consists of 60 protein copies forming a ∼25‐nm particle, whereas HPV is even larger, comprising 360 identical L1 proteins arranged into a ∼60‐nm icosahedral shell. Simulating assemblies of this scale at AA resolution would require millions of atoms and exceptionally long trajectories to achieve equilibration, far exceeding accessible computational resources. Consequently, most existing computational studies rely either on HCG models or on small AA subunits, leaving no practical way to connect mesoscopic CG dynamics to atomistic detail. To bridge this gap, we integrate our FLCG (ESI Figure S3) approach^8^ and progressive backmapping (ESI Figure S4) with the ProNet Backmapping framework, leveraging the neural‐network‐guided reconstruction to hierarchically convert models across resolutions. In this section, integrating the FLCG method, progressive backmapping method^8^ and ProNet Backmapping, we coarse‐grained the AAV2 and HPV capsids from their AA models to 1‐residue/site and 3‐residue/site CG representations, then progressively backmapped the 3‐residue/site models to 1‐residue/site and finally to full AA coordinates with ProNet Backmapping. This workflow enables, for the first time, accurate and efficient backmapping of entire viral assemblies from HCG models to atomistic resolution.

The overall workflow is summarized in Figure 5A. For each protein within the AAV structure, a CG representation was first generated by mapping the AA model into the 1-residue/site CG model by applying the center of mass scheme for each residue. Subsequently, three neighboring beads were grouped into one single bead by using the FLCG method (ESI Figure S3) to construct a 3-residue/site model as illustrated in Figure 5B. The ProNet Backmapping model then progressively backmapping the protein structure from 3-residue/site into 1-residue/site, and ultimately to the AA model. For the AAV system, the AA models of the 60 constituent proteins were predicted individually and then reassembled according to their spatial arrangement. Figure 5D presents the complete coarse-graining and backmapping process for AAV. The initial structure is obtained from X-ray crystallography, followed by progressively CG models and backmapped AA models. The predicted AA model exhibits an overall RMSD of approximately 3.5 Å relative to the X-ray structure, where the RMSD of the backbone is reduced to below 2.5 Å, as shown in Figure 5C.

**Figure 5.**
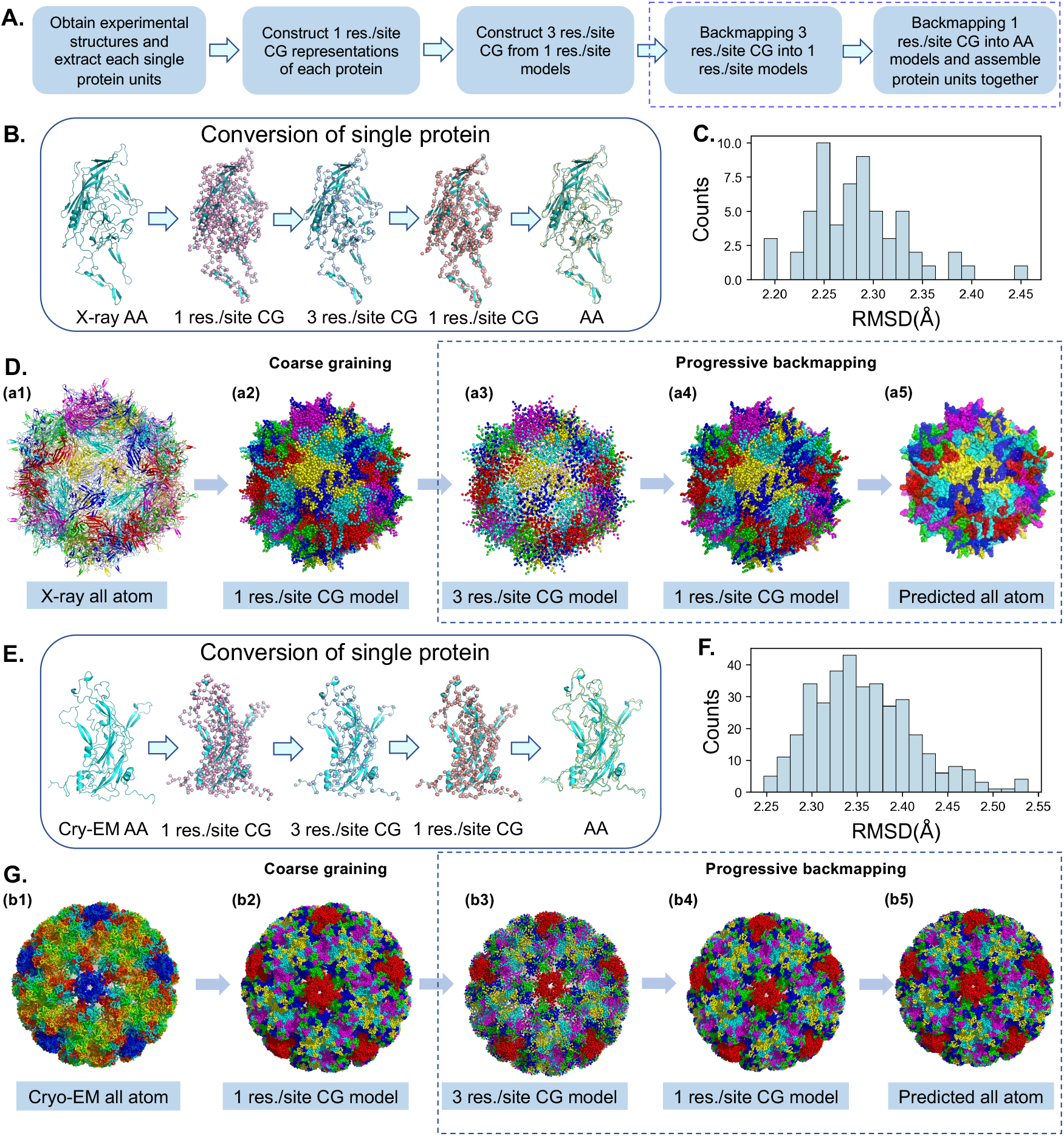
(A) Workflow for transforming AAV2 and HPV models across multiple resolutions. (B) Coarse-graining and backmapping of a single AA2 protein. (C) Distribution of RMSD values for predicted all-atom AA2 models. (D) AAV2 models at different resolutions: (a1) X-ray obtained all-atom structure, (a2) coarse-grained 1-residue/site CG model, (a3) coarse-grained 3-residue/site model, (a4) backmapped 1-residue/site model, (a5) backmapped all-atom model. (E) Coarse-graining and backmapping of a single HPV protein. (F) Distribution of RMSD for predicted all-atom HPV models. (G) HPV models at different resolutions: (b1) Cryo-EM obtained all-atom structure, (b2) coarse-grained 1-residue/site CG model, (b3) coarse-grained 3-residue/site model, (b4) back-mapped 1 res./site model, (b5) back-mapped all-atom model.

This scheme was further applied to the HPV structure, which consists of 360 proteins, with each six of them representing a hexamer, resulting in an overall hollow icosahedral architecture. The results of the coarse-graining and backmapping process are shown in Figure 5G. The reconstructed structure achieves an overall RMSD of approximately 3.5 Å with the backbone RMSD being even lower, falling below 2.5 Å as presented in Figure 5G. These values indicate a high level of structural accuracy, demonstrating the method’s ability to preserve both global geometry and local atomic details in large, symmetrical assemblies.

### Progressive backmapping to test protein mutations

Mutations are commonly introduced into proteins to enhance their performance or confer new functions, such as improving delivery efficiency, increasing binding affinity, or enhancing overall structural stability.^72–75^ Our ProNet Backmapping approach is useful not only for modeling wildtype proteins but also for testing various mutations or variants. To demonstrate this application, we explore a case study of the AAV2 pentamer, as mutations in AAV vectors are often created to improve gene therapy effectiveness by enhancing targeting precision, evading immune detection, and increasing transgene expression. Specifically, we introduced mut19-mut25, each of which has 5 residues (Figure 6A), into the AAV2 model during the backmapping process. Previous work has shown that these mutations influence AAV2 stability and assembly.^76^ Using a 1-residue/site CG model built from the wildtype AA structure, we backmapped both the wildtype and mutant AA models with ProNet Backmapping. Although the CG models of the wildtype and mutant are identical, the AA models and subsequent MD simulations clearly differentiate them: the RMSD of the mutant pentamer increases significantly to 10 Å within 20 ns, whereas the wildtype model remains mostly below 6 Å (Figure 6E). Although the mutant does not dissociate, there is a trend of increasing subunit distances (Figure 6B); residues within each subunit tend to switch neighboring contacts more frequently, as also shown by the contact maps in ESI Figure S6 and S7. Further RMSF analysis reveals that the mutant is overall more dynamic than the wildtype, especially in the N- and C-termini (Figure 6F). We can verify that these mutations significantly influence the stability of the AAV2 assembly using backmapping, in good agreement with previous experimental findings.^76^ Because the mutation region in our model is exclusive to the N-terminus, the observation of a high RMSF in the C-terminus demonstrates a global effect captured by our backmapped AA model and simulation. Given the efficiency of our approach, it is possible to screen or test a wide range of mutations and select desirable sequences and properties using backmapping. In general, our backmapping framework can serve as an efficient tool for identifying engineerable hotspots in AAV and similar vectors for drug delivery applications.

**Figure 6.**
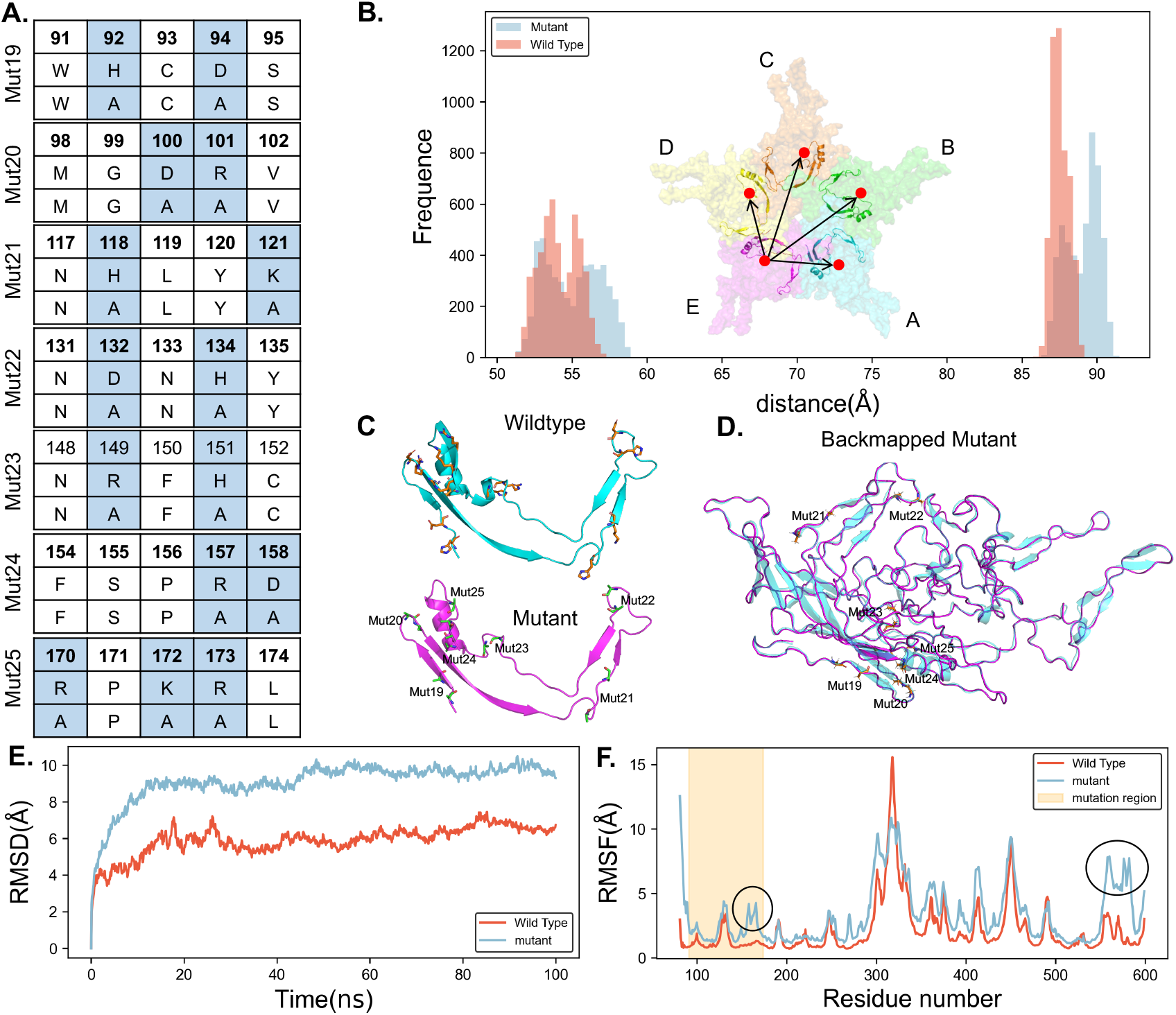
(A) The highlighted mutation introduced into the AAV2 pentamer. (B) Radial distribution function between the 5 subunits of the pentamer. The carton of the insertion graph highlights mutant region. (C) The mutated regions of the protein corresponded to the region in the wild type; This segment spans from residue 92 to 173 and corresponds to region has mut19-25. (D) The backmapped mutant (magenta) of a subunit from ProNet Backmapping, backmapped mutant residues are shown as sticks. The backmapped single subunit structure(magenta) algins well with the reference mutant structure (cyan), with a backbone RMSD of approximately 1.2 Å. (E) RMSD and (F) RMSF for both the wild-type and mutant pentamers.

## Conclusion

In this work, we introduce a progressive backmapping framework which can stepwise reconstructing accurate AA protein structures from HCG models. This approach integrates ProNet Backmapping, a neural‐network–enabled, thermodynamically consistent model, the FLCG coarse graining approach, and our previous advances in progressive resolution refinement. Using this framework, we demonstrate—for the first time—the ability to backmap entire viral assemblies such as AAV2 and HPV VLPs from 3-residue/site to 1-residue/site representations, then to full AA resolution. Across diverse proteins, ProNet Backmapping consistently achieves sub‐2 Å accuracy, with particularly strong performance in backbone reconstruction, flexible linkers, and multidomain architectures. The method also preserves residue‐level dynamics and thermodynamic consistency, overcoming long‐standing limitations of deterministic or rule‐based approaches.

Beyond validating backmapping for large protein assemblies, this work establishes a backmapping framework as a scalable tool for bridging atomistic and mesoscopic simulations, enabling new research directions that were previously inaccessible due to computational constraints. Its generality paves the way for future applications such as reconstructing AA detail from long‐timescale CG simulations of viral capsids, nanoparticles, and protein assemblies; refining intrinsically disordered regions; integrating with enhanced‐sampling or AI‐driven MD workflows; interpreting intermediate‐resolution cryo‐EM maps; and supporting large‐scale mutational design for engineered biomolecular systems. Together, these capabilities position backmapping as a versatile and powerful framework for advancing multiscale modeling, structural biology, and nanomedicine.

## Supporting information

ESI

## Acknowledgements

The authors thank Dr. Jonathon B. Ferrell for helpful discussions. This work was supported by a National Science Foundation Career award (CHE-2410514) to J.L. S.T.S. was supported by an NIH R35 Award (GM147579).

## Supporting information

Workflow for pre-processing input data; list of PDB structures; RSMF of MDPs; fixed-length coarse-graining; Progressive backmapping; Backmapping of small proteins; Simulation of AAV2 pentamer; Contact maps of pentamers.

