## Supplementary material for "Progressive Backmapping of Highly Coarse-Grained Protein Models": ESI

**Electronic Supplementary Information:**

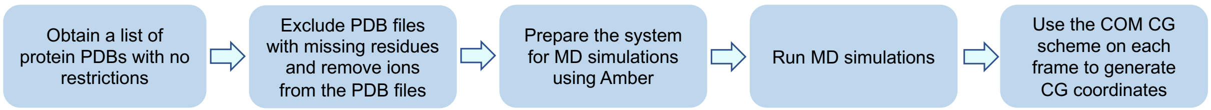


**Figure S1.** Cartoon depicting the process of preparing input data.

**Table S1.** All proteins used for both training and validation in this study.

| | | | | | | | | | |
|---|---|---|---|---|---|---|---|---|---|
| 12E8 | 1A3H | 1A4U | 1A7U | 1A88 | 1AGJ | 1AKO | 1ARL | 1AUO | 1BEC |
| 1BHE | 1BKP | 1BQC | 1BUE | 1BYI | 1CNS | 1CZ1 | 1DIX | 1DJA | 1DQ0 |
| 1DUZ | 1EDG | 1EQP | 1ERZ | 1F00 | 1F5Z | 1FBA | 1FSF | 1FTR | 1G24 |
| 1G8A | 1GQN | 1GVL | 1GZJ | 1H6U | 1HRD | 1HTR | 1HXH | 1HYL | 1I7U |
| 1IDK | 1IJB | 1IU8 | 1JFL | 1JGV | 1JLN | 1KCV | 1KS9 | 1L7A | 1L8F |
| 1LBV | 1LP8 | 1LU9 | 1M5S | 1M6O | 1MIZ | 1MMI | 1N7P | 1NBZ | 1NGZ |
| 1NLB | 1OA4 | 1OCK | 1OJQ | 1OMP | 1ONR | 1ORS | 1P3C | 1PE9 | 1PIP |
| 1PP3 | 1PXZ | 1PZ5 | 1QCX | 1QTS | 1RC9 | 1RL0 | 1T06 | 1T4D | 1THV |
| 1TIB | 1TJE | 1TUX | 1U00 | 1UGH | 1USG | 1VAX | 1X1E | 1XDW | 1XH3 |
| 1XSZ | 1XVM | 1Y6I | 1YPI | 1YT4 | 1YUO | 1Z15 | 1ZAH | 1ZHL | 1ZSD |
| 1ZZG | 2A6Z | 2AJU | 2B5R | 2BAA | 2BKR | 2BNU | 2CB5 | 2CGA | 2CLZ |
| 2CV3 | 2CYG | 2D0I | 2D5J | 2DUC | 2ERF | 2EXO | 2EYI | 2FAT | 2FB4 |
| 2FZ3 | 2G2U | 2G5X | 2GAS | 2GGO | 2H2Z | 2H6P | 2HAD | 2HJK | 2HLC |
| 2HOB | 2HVM | 2IE8 | 2LAO | 2NW3 | 2O6S | 2OF3 | 2OKT | 2OMZ | 2OR2 |
| 2PA6 | 2PET | 2QCY | 2QHT | 2SIL | 2V2W | 2VLL | 2VU8 | 2WBZ | 2WPL |
| 2X8X | 2XFX | 2YMU | 2YNO | 2YPK | 2YXP | 3AAP | 3APP | 3BWA | 3CTK |
| 3D25 | 3ER5 | 3F7M | 3GSW | 3H6J | 3ILS | 3KLA | 3KPQ | 3LIG | 3LKO |
| 3LN5 | 3LY3 | 3M4D | 3M66 | 3N4I | 3OT7 | 3OXR | 3P8T | 3PTL | 3PWU |
| 3RP2 | 3RRS | 3SEB | 3TUA | 3UTQ | 3VFS | 3W3E | 3W4Q | 3WA1 | 3WP5 |
| 3WY8 | 3X13 | 4AXU | 4CFI | 4DIY | 4DJ5 | 4FCU | 4G5Z | 4G6K | 4GKU |
| 4GUZ | 4H20 | 4I4N | 4J4R | 4JCN | 4JJO | 4JZC | 4KEL | 4L9R | 4LE8 |
| 4LSW | 4MCK | 4MHP | 4NT6 | 4O2C | 4OZX | 4P9N | 4PBO | 4PMH | 4PR5 |

| | | | | | | | | | |
|---|---|---|---|---|---|---|---|---|---|
| 4R1N | 4RVS | 4RXV | 4TX7 | 4V38 | 4WJS | 4WUM | 4X7S | 4XIO | 4YHE |
| 4ZPB | 4ZTP | 5A8U | 5AMZ | 5AVG | 5C0D | 5DDT | 5DJ7 | 5DK1 | 5DTX |
| 5DZ9 | 5E5B | 5EFS | 5EO1 | 5EWT | 5GLX | 5GN2 | 5GQP | 5GS7 | 5GY3 |
| 5H0Q | 5H28 | 5H4E | 5H5Z | 5HHP | 5IB1 | 5IHW | 5J2V | 5KEH | 5KJV |
| 5MAL | 5ORI | 5OYX | 5OZ9 | 5P00 | 5P11 | 5P20 | 5P31 | 5P42 | 5P53 |
| 5P64 | 5P75 | 5P86 | 5TUN | 5U3P | 5UCB | 5VTL | 5VWH | 5XCY | 5XOS |
| 5Y33 | 5YMW | 5YNS | 5YPV | 5YR3 | 5YSX | 5YUP | 5Z2D | 5ZFH | 6A41 |
| 6AFM | 6CL7 | 6D0A | 6D29 | 6E3D | 6E4D | 6F4M | 6FAB | 6GFV | 6GGP |
| 6HY2 | 6I3B | 6IAS | 6IM4 | 6J1W | 6JTP | 6KGJ | 6L8T | 6NZS | 6OKJ |
| 6PKP | 6PYW | 6PZ5 | 6QE3 | 6VT3 | 6WTM | 6XIA | 6Y2E | 7AHL | |

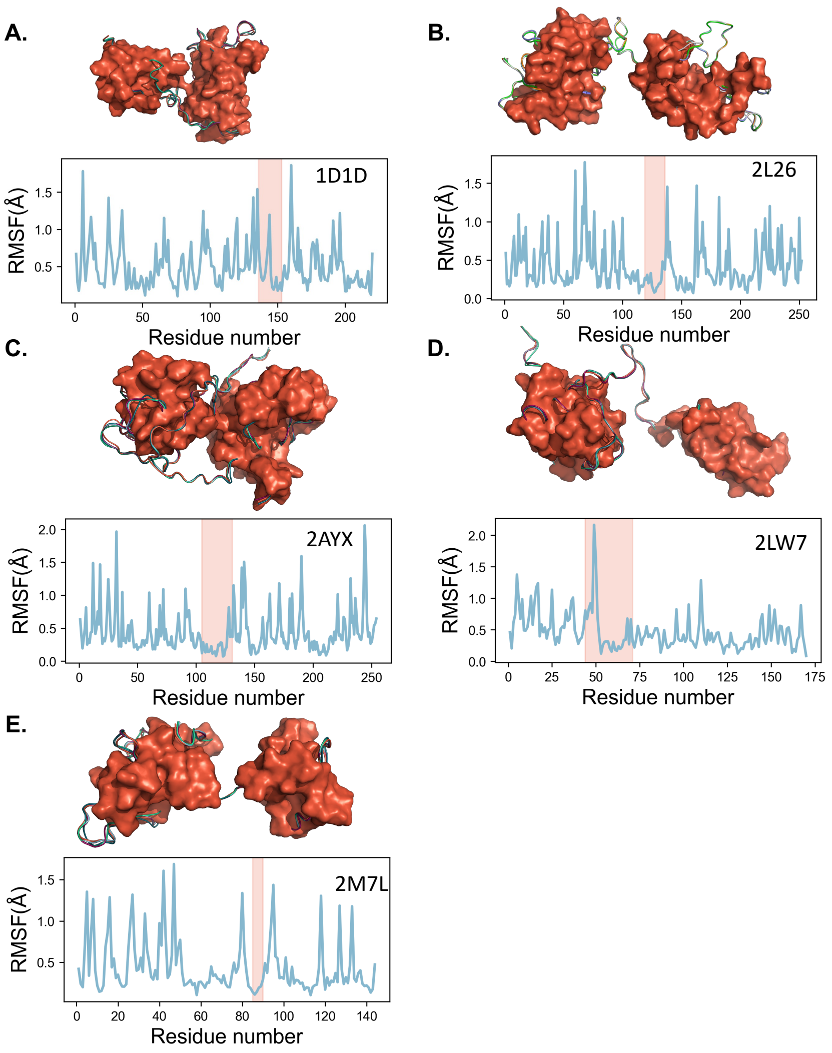


**Figure S2.** RMSF of example multidomain proteins, and their corresponding snapshots, with the linker region highlighted in pink.

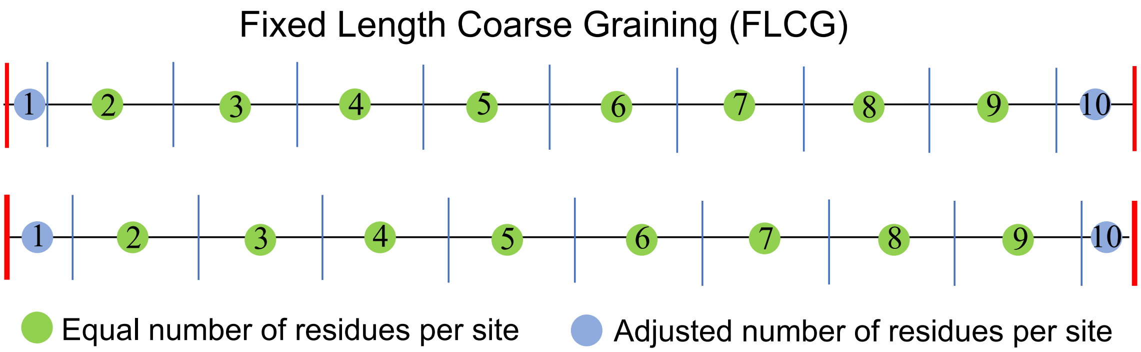


**Figure S3.** Schematic diagram illustrating two possible CG assignments from FLCG output of 10-site HCG models. Green circles show the CG sites of an equal number of residues. The termina CG sites (blue circles) can have fewer residues. An optimal assignment is to keep both terminal CG sites with a similar number of residues.

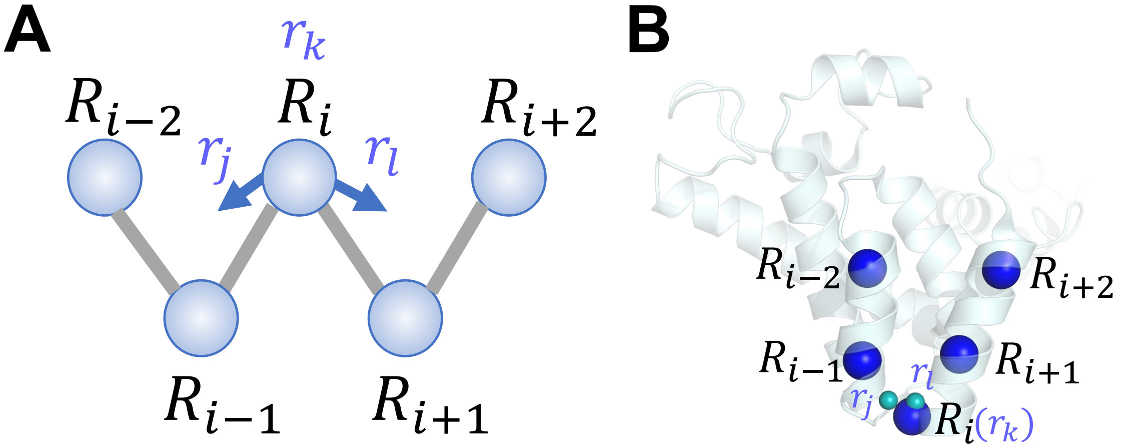


**Figure S4.** (A)Simple geometry-based rules in progressive backmapping from lower-resolution 3 res./bead CG coordinates R to higher-resolution 1 res./bead coordinates r. (B) A small portion of protein (PDB ID: 1D1D) illustrates the method, where the blue beads belong to the lower-resolution model and cyan beads belong to the higher-resolution model. The sphere of $R_i$ and $r_k$ overlap in this model.

**Table S2.** Several additional backmapped proteins with fewer than 100 residues.

| PDID | Overall (Å) | Backbone (Å) | Sidechain (Å) |
|---|---|---|---|
| 1CHL | 2.98 | 1.78 | 3.3 |
| 1F81 | 3.92 | 2.74 | 4.23 |
| 1DEF | 3.55 | 2.09 | 3.85 |
| 1AGT | 3.14 | 2.36 | 3.36 |
| 1C55 | 3.5 | 2.3 | 2.83 |
| 1DME | 3.13 | 2.05 | 3.49 |
| 1CCQ | 3.64 | 2.35 | 3.97 |
| 1CCN | 3.49 | 2.3 | 3.86 |
| 1CB9 | 3.02 | 2.14 | 3.26 |
| 1CAD | 3.08 | 2.18 | 3.35 |
| 1EHS | 4.0 | 2.8 | 4.36 |
| 1ED0 | 3.4 | 2.27 | 3.75 |
| 1DTV | 3.18 | 2.1 | 3.49 |
| 1DEC | 3.4 | 2.05 | 3.77 |
| 1D1H | 3.1 | 2.18 | 3.34 |
| 1ACW | 3.11 | 2.57 | 3.67 |
| 1AOO | 3.17 | 2.3 | 3.45 |
| 1APF | 3.58 | 2.46 | 4.21 |
| 1ARD | 3.77 | 2.66 | 4.01 |
| 1ARE | 3.99 | 2.7 | 4.25 |
| 1B7B | 3.47 | 2.29 | 3.79 |
| 1B8W | 3.53 | 2.08 | 3.89 |
| 1B9G | 3.7 | 2.4 | 4.04 |

**All-atom MD Simulations of AAV2 pentamer:**

The simulation systems were created using the Solution Builder model in CHARMM-GUI[1,2]. The force field used in the simulations was CHARMM36m[3] with the TIP3P water model[4]. A solvation box measuring 23.0 nm on each side was utilized to house the backmapped AAV pentamer complex (original and mutated), ensuring ample room for any necessary deformations. Two systems were neutralized by adding counterions. The MD simulations were performed using the GROMACS2024.3 software package[5,6]. All systems were energy-minimized through a maximum of 50000 steps to eliminate non-physical contacts and interactions.

Subsequently, an NPT ensemble with a 1 ns time step was used to equilibrate the systems. The LINCS algorithm was used to constrain bond lengths between heavy and hydrogen atoms[7]. Simulations were performed under a constant temperature of 303.15 K and a constant pressure of 1 atm using the Parrinello-Rahman method[8]. Periodic boundary conditions were applied in all directions, and the Particle mesh Ewald (PME) method was used to compute electrostatic interaction.[9] The cut-off distance of van der Waals interaction and Coulomb interaction were both set to 12 Å. The duration of production simulations was 100ns. Analysis of simulation results was done using GROMACS2024.3.[10] Simulation results were visualized using UCSF Chimera and Pymol.

A.

B.

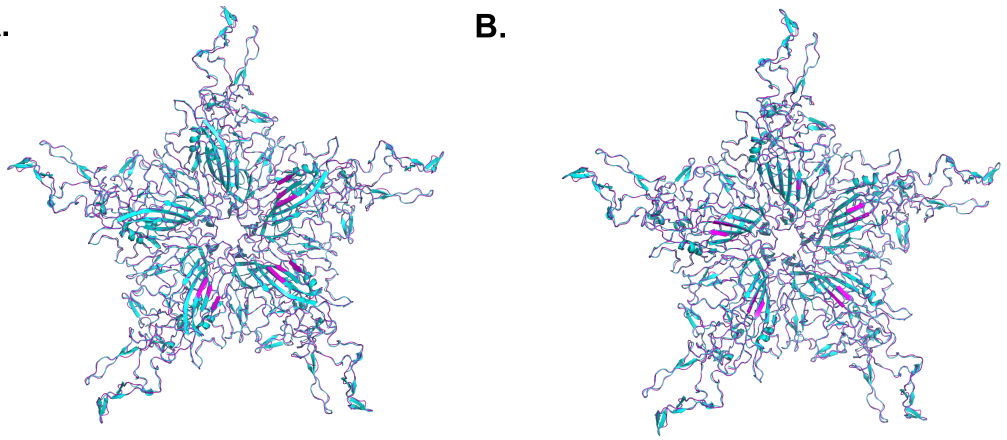

**Figure S5.** (A)Wild type AAV2 pentamer (cyan) and its corresponding backmapped structure (magenta) derived from 1 res./site CG model. (B) Mutant AAV2 pentamer (cyan) and its corresponding backmapped structure (magenta) derived from 1 res./site CG model.

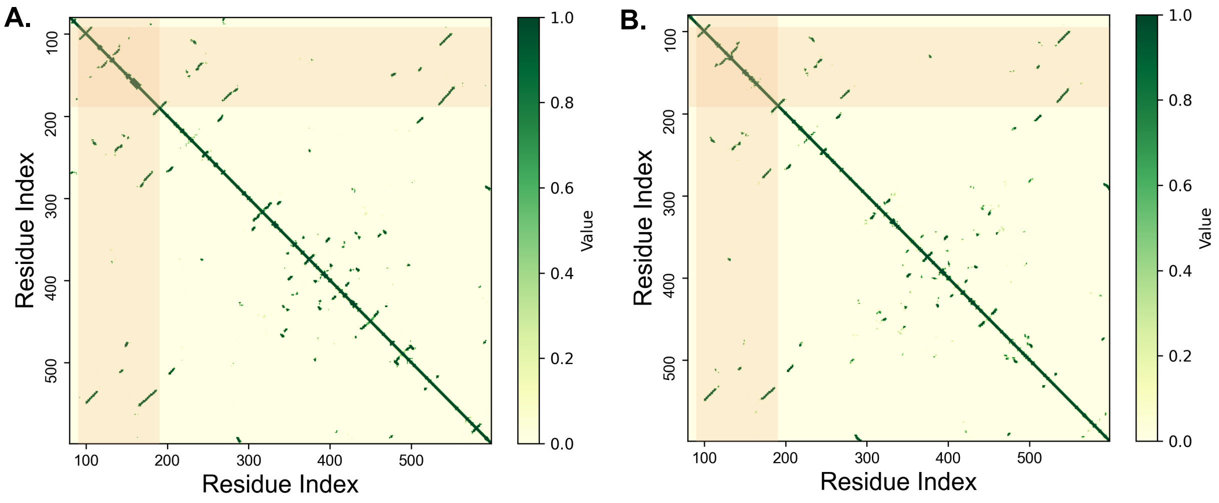


**Figure S6.** (A) Contact map of subunit A in the wild type pentamer. (B) Contact map of subunit A in the mutant pentamer. The highlight region indicates the site of the introduced mutation.

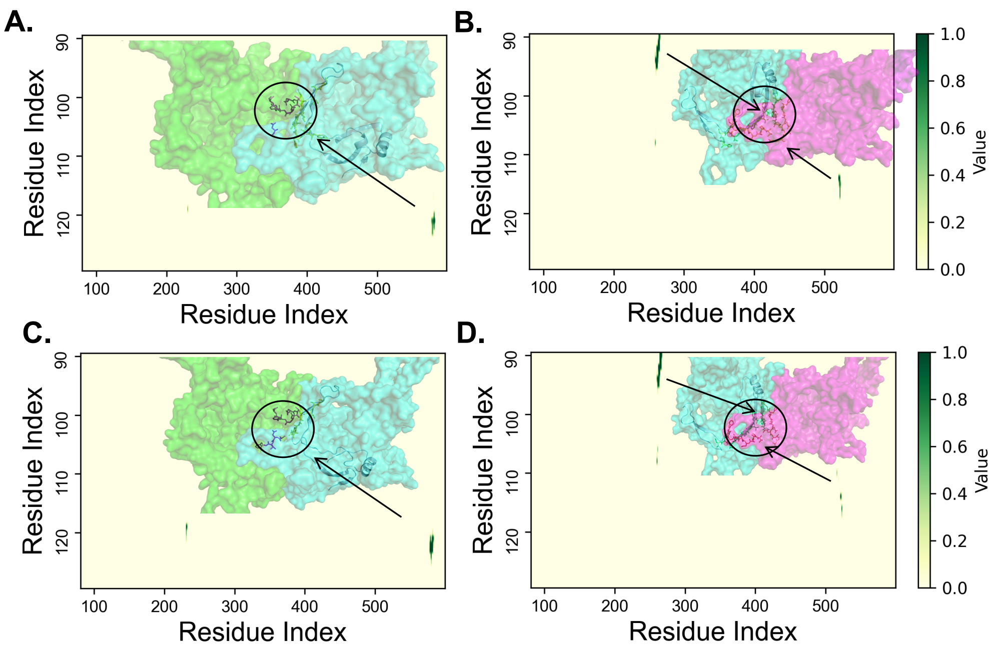


**Figure S7.** Intermolecular contact maps between neighboring subunits: wild type protein A-B (A), A-E(B), mutant A-B (C), and A-E (D).
